# LEXAS: a web application for life science experiment search and suggestion

**DOI:** 10.1101/2021.12.05.471323

**Authors:** Kei K Ito, Yoshimasa Tsuruoka, Daiju Kitagawa

## Abstract

**Motivation:** In cellular biology, researchers design wet experiments by reading the relevant articles and considering the described experiments and results. Today, researchers spend a long time exploring the literature in order to plan experiments.

**Results:** To accelerate experiment planning, we have developed a web application named LEXAS (Life-science EXperiment seArch and Suggestion). LEXAS curates the description of biomedical experiments and suggests the experiments on genes that could be performed next. To develop LEXAS, we first retrieved the descriptions of experiments from full-text biomedical articles archived in PubMed Central. Using these retrieved experiments and biomedical knowledgebases and databases, we trained a machine learning model that suggests the next experiments. This model can suggest not only reasonable genes but also novel genes as targets for the next experiment as long as they share some critical features with the gene of interest.

**Availability and implementation:** LEXAS is available at https://lexas.f.u-tokyo.ac.jp/ and provides users with two interfaces: search and suggestion. The search interface allows users to find a comprehensive list of experiment descriptions, and the suggestion interface allows users to find a list of genes that could be analyzed along with possible experiment methods. The source code is available at https://github.com/lexas-f-utokyo/lexas.

**Supplementary information:** Supplementary data are available at Bioinformatics online.

## 1 Introduction

In cellular biology, researchers often face questions like “Which gene should we target in the next experiment? “ and “What experiment should we perform for that gene?” once they finish an experiment on a gene. They usually address such questions by consulting the related literature and biomedical databases. Due to the large body of literature and many databases available today, researchers spend a long time planning their experiments.

There are many text-mining-based web applications that can help researchers quickly grasp various gene-related information described in the literature (Tsuruoka et al., 2008; Shen et al., 2018). For example, STRING (Szklarczyk et al., 2021) can generate a relationship network diagram with the target gene at the center based on text-mining and several gene features like expression levels. However, none of the existing systems can directly answer the aforementioned questions, i.e., they do not tell users which genes should be targeted or what experiment method should be performed in the next experiment.

To help researchers efficiently plan experiments on genes, we have developed a web application named LEXAS (Life-science EXperiment seArch and Suggestion). Using the information about the gene analyzed in the previous experiment, LEXAS can suggest genes to be investigated in the next experiment along with possible experiment methods. LEXAS can also be used as an accurate search engine for biomedical experiments described in the literature.

## 2 Methods and results

This section describes the text data and machine learning models used in LEXAS.

### 2.1 Text preprocessing

We collected sentences that describe gene-related experiments from the biomedical literature. First, we downloaded all full-text articles archived in PubMed Central (PMC) and extracted only the result sections. Then, we extracted the sentences that contain at least one gene name and one experiment method. As a result, 34,586,978 sentences were retrieved from 1,068,342 articles.

### 2.2 Relation extraction for experiment retrieval

To obtain information about the experiments performed on genes from the collected sentences, we built a relation extraction model using bio-BERT (Lee et al., 2020). First, we created a dataset by manually annotating 1658 sentences for gene-experiment relations. Here, a gene and an experiment are defined to be in a gene-experiment relation if the experiment was performed on the gene. We then masked a gene name and an experiment method in each sentence with special tokens, [GENE] and [EXPE], respectively. Finally, the bio-BERT model was finetuned to predict whether [GENE] and [EXPE] are in a gene-experiment relation.

The dataset was divided into a training set and a test set, and the precision of the relation extraction task on the test set was 0.824, with a relative recall of 0.810. As a baseline, a precision of 0.271 and a relative recall of 1.0 can be obtained by considering any combination of a gene name and a method in each sentence to be in the gene-experiment relation.

Using this model, we extracted 9,689,108 pairs of a gene and an experiment that represent gene-experiment relations.

### 2.3 Predicting genes for the next experiment

Using the above-mentioned data of gene-experiment relations, we trained a machine learning model for experiment suggestion. In this work, we assume that the order in which experiments are mentioned in text reflects the actual order of the performed experiments. Therefore, the task of the model is predicting the next target gene and experiment method using information about the previous experiment.

Several major biomedical knowledgebases and databases were used to construct feature vectors representing the context of the experiments. These information sources associate each gene with some features. For example, the Gene Ontology (Carbon et al., 2019) assigns several terms to each gene that describe biological processes, subcellular localization, or signal pathways. Using the features of target genes in two consecutive experiments, we constructed feature vectors and then trained an XGBoost model (Chen and Guestrin, 2016). The model was trained using the experiments described in articles from 2010 to 2019 and evaluated with the experiments described in 2020.

To evaluate the accuracy of the suggested target genes, we calculated the mean average precision (MAP@100) score. For each gene mentioned in the articles published in 2020, the model was used to calculate the probabilities of occurrence for all human genes as the target of the next experiment. We then ranked the genes by their probabilities and obtained a MAP score of 0.369. This score is significantly better (p < 0.0001, bootstrapping method) than a baseline MAP score of 0.310 that can be obtained using the “scored links between proteins” provided by the STRING database. In particular, for the genes not studied before 2019, the MAP score of LEXAS was 0.278 whereas that of STRING was 0.109, suggesting that our system can predict reasonable genes even if the query gene has not been studied yet.

For the suggestion of experiment methods, we categorized the experiment methods into 19 groups and trained a multi-class logistic regression classifier to predict the experiment category in the next experiment. The accuracy of method suggestion was 0.452.

## 3 User interface

LEXAS provides users with two interfaces: search and suggestion.

### 3.1 LEXAS Search

The search interface allows users to find a comprehensive list of experiment descriptions extracted by the relation extraction model presented in Section 2.2. Given a gene name and the category of an experiment method by the user, the system shows the list of matching experiment descriptions as a table. The table shows the sentences that describe the experiment, along with the confidence score output by the relation extraction model.

### 3.2 LEXAS Suggestion

The suggestion interface allows users to find a list of genes that could be analyzed after an experiment on a given gene. The important features are shown along with their SHAP values (Lundberg and Lee, 2017), helping the user understand why those genes are suggested.

The system also allows users to choose between two machine learning models that are used for the suggestion. One is a “reliable” model that uses seven major biomedical databases such as the BioGRID and four knowledgebases such as the Gene Ontology, which we evaluated in Section 2.3. This model is suitable for those who are seeking reliable suggestions in line with a published body of knowledge. By contrast, the other “exploratory” model is built using biomedical databases alone. This model is thus suitable for those who are seeking novel connections based on relatively objective features of genes.

## 4 Conclusion

To help cellular biologists design experiments, we have developed a web application named LEXAS that curates the descriptions of performed experiments from the literature and suggests the experiments that could be performed next. LEXAS is available at https://lexas.f.u-tokyo.ac.jp.

**Figure 1.**
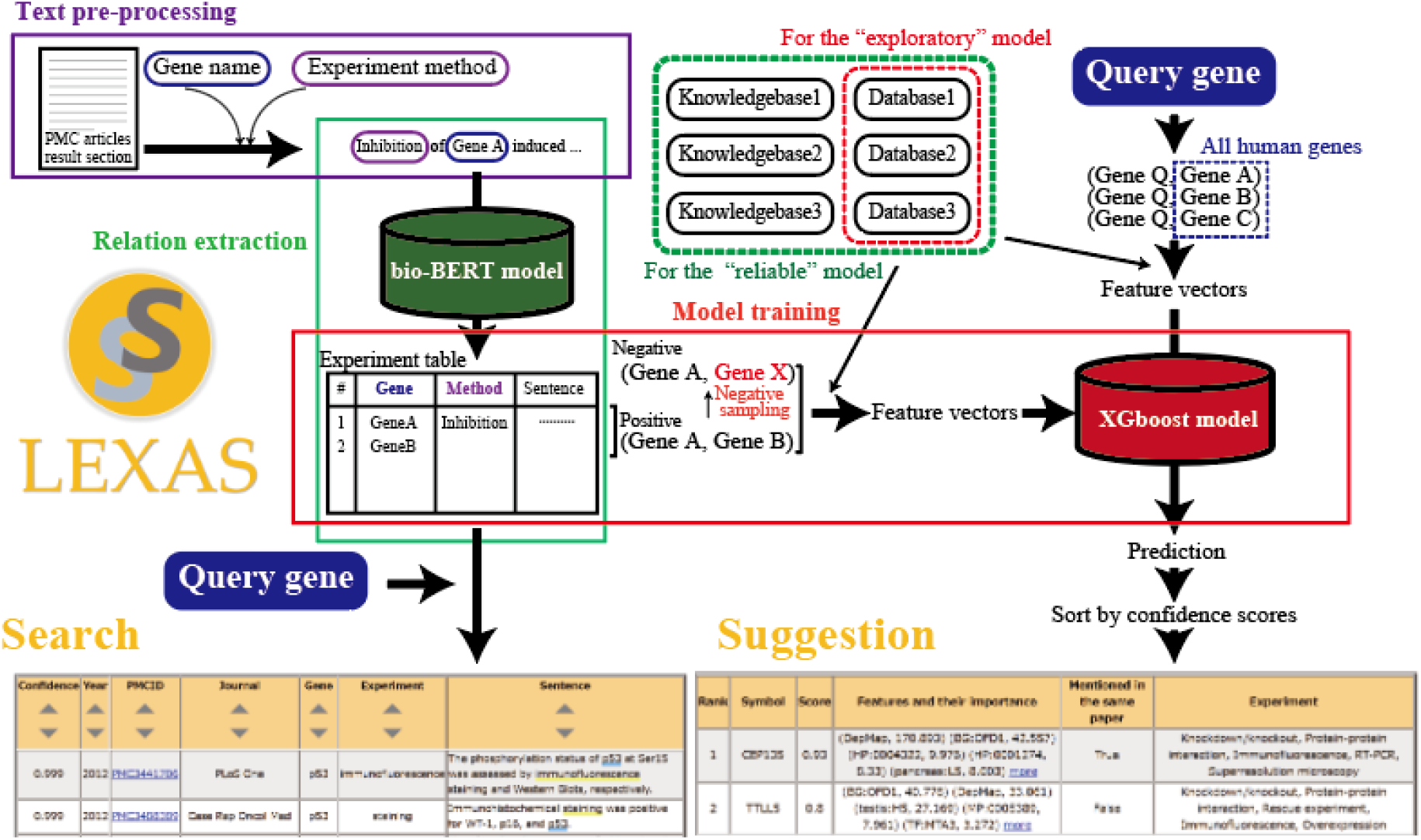
Overview of LEXAS. The result tables of LEXAS Search and Suggestion can be obtained when entering “TP53, immunofluorescence” and “Cep63”, respectively.

## Acknowledgments

We gratefully acknowledge T. Mizuno, N. Kono, Y. Kishi, and Kitagawa lab members for constructive feedback on the user interface of LEXAS.

## Funding

This work has been supported by a grant-in-aid for scientific research (S, 19H05651) and the IIW program of The University of Tokyo.

## Conflict of Interest

**none declared**.

## Notes

https://lexas.f.u-tokyo.ac.jp/

### Competing Interest Statement

The authors have declared no competing interest.

## References

Carbon, S. et al. (2019) The Gene Ontology Resource: 20 years and still GOing strong. Nucleic Acids Res., 47, D330–D338.

Chen, T. and Guestrin, C. (2016) XGBoost: A scalable tree boosting system. In, Proceedings of the ACM SIGKDD International Conference on Knowledge Discovery and Data Mining.

Lee, J. et al. (2020) BioBERT: A pre-trained biomedical language representation model for biomedical text mining. Bioinformatics.

Lundberg, S.M. and Lee, S.I. (2017) A unified approach to interpreting model predictions. In, Advances in Neural Information Processing Systems.

Shen, J. et al. (2018) DLAD4U: deriving and prioritizing disease lists from PubMed literature. BMC Bioinformatics, 19, 495.

Szklarczyk, D. et al. (2021) The STRING database in 2021: Customizable protein-protein networks, and functional characterization of user-uploaded gene/measurement sets. Nucleic Acids Res.

Tsuruoka, Y. et al. (2008) FACTA: a text search engine for finding associated biomedical concepts. Bioinformatics, 24, 2559–2560.

